# Is cathodal prefrontal transcranial direct current stimulation capable of affecting inhibitory control and sustained attention? A single-blind, crossover, sham-controlled study

**DOI:** 10.1101/2020.01.20.912287

**Authors:** Soraya Lage de Sá Canabarro, Cássia Karolina Paniago, Priscilla Magalhães Santos, Lorena da Silva Rosa, Vitória Espíndola Leite Borges, Daniel Pimentel McManus, Ana Garcia, Corina Satler, Joaquim P. Brasil-Neto, Maria Clotilde Henriques Tavares

## Abstract

**Background:** Anodal transcranial direct current stimulation (a-tDCS) has been shown to promote performance improvement of normal individuals in tests of executive function, including tasks that demand sustained attention and inhibitory control. The presumed mechanism is facilitation of prefrontal cortex activation, since a-tDCS is thought to increase cortical excitability. Only a few studies, however, have investigated the effects of inhibitory, cathodal tDCS (c-tDCS) on cognitive tasks, and reported results are often inconsistent. Studies about the effects of c-tDCS upon accuracy and reaction times are particularly scant.

**Objective/Hypothesis:** This study assessed the effects of inhibitory c-tDCS over the left dorsolateral prefrontal cortex (l-DLPFC) on the performance of neurologically intact young adults in Stroop and reaction time tests.

**Methods:** Seventeen healthy undergraduate students (ten women) performed Stroop and reaction time tasks after delivery of c-tDCS (1 mA, 20 min) over l-DLFPC or a sham session. All subjects underwent both real and sham sessions, which were separated by an interval of one week. We hypothesized that c-tDCS might lead to an impairment of inhibitory control and attention abilities.

**Results:** We found an interference effect on the Stroop task and also a ceiling effect on the reaction Time task. There were no statistically significant performance differences in any of the neuropsychological tests as a function of stimulation condition and/or subject gender.

**Conclusions:** C-tDCS over the l-DLPFC of neurologically intact young individuals did not affect performance in Stroop Test accuracy or in reaction times, irrespective of subject gender. These results raise the possibility that c-tDCS inhibitory effects, well documented for the primary motor area, do not necessarily apply to higher order associative areas. The assumption that c-tDCS has inhibitory effects upon any cortical area, common in clinical trials, should be made with caution.

## Introduction

Transcranial direct current stimulation (tDCS) has been extensively studied as a means of modulating cortical excitability in both neurologically intact subjects and in patients with neuropsychiatric disorders (Nitsche and Paulus 2000; Nitsche, Cohen, Wassermann, et al. 2008; Boggio, Bermpohl, Vergara, et al. 2007; Jacobson, Javitt, and Lavidor 2011; Miniussi and Ruzzoli 2012; Paulus, Antal, and Nitsche 2013). Regarding the motor cortex, depolarization effects upon neuronal cell membrane, which induce behavioral facilitation, have been described after anodal stimulation (a-tDCS), while opposite effects have been observed during cathodal stimulation (c-tDCS) (Paulus, Antal, and Nitsche 2013; Nitsche and Paulus 2000). In general, however, tDCS effects depend on technical parameters used during stimulation (current density, stimulation duration, positioning and size of the electrodes) as well as on the interaction between these parameters and subject-related variables, such as gender, age, and cognitive state (Miniussi and Ruzzoli 2012; Plewnia, Zwissler, Längst, et al. 2013).

Recent studies have demonstrated that prefrontal a-tDCS is able to promote improvement in the performance of normal individuals in tests of executive functions (Sarkis, Kaur, and Camprodon 2014), including tasks that demand sustained attention or vigilance (Nelson, McKinley, Golob, et al. 2014) and inhibitory control, such as the Stroop task (Jeon and Han 2012), Stop Signal task (Hsu, Tseng, Yu, et al. 2011; Jacobson, Javitt, and Lavidor 2011; Ditye, Jacobson, Walsh, et al. 2012; Kwon and Kwon 2013), Go-no-go task (Boggio, Bermpohl, Vergara, et al. 2007; Beeli, Casutt, Baumgartner, et al. 2008) and Choice Reaction Time task (Kang, Kim, and Paik 2012).

The effects of c-tDCS over the prefrontal cortex on inhibitory control and attention mechanisms, however, have not been extensively studied (Sarkis, Kaur, and Camprodon 2014), with a few exceptions (Nelson, McKinley, Golob, et al. 2014; Jacobson, Javitt, and Lavidor 2011; Ditye, Jacobson, Walsh, et al. 2012). Moreover, a recent detailed systematic review of published a-tDCS and c-tDCS studies (Dedoncker, Brunoni, Baeken, and Vanderhasselt 2016) found no effect of c-tDCS on reaction time or accuracy during performance of neuropsychological tests.

Reasons for such lack of effect of c-tDCS are unclear. The assumption that a-tDCS has excitatory effects whereas c-tDCS is inhibitory is based on studies on the motor cortex (Nitsche and Paulus 2000) and may a be flawed one. The effects of tDCS upon a tertiary associative cortical area such as the prefrontal cortex might be different from those obtained in the primary motor cortex (Dedoncker, Brunoni, Baeken, and Vanderhasselt 2016; Jacobson, Javitt, and Lavidor 2011). Genetic variability in response to c-tDCS may be another explanation for the fact that a few studies show clear-cut effects of c-tCDS upon executive functions (Filmer, Mattingley, and Dux 2013; Leite, Carvalho, Fregni, and Gonçalves 2011), whereas most trials have shown a lack of effect (Dedoncker, Brunoni, Baeken, and Vanderhasselt 2016). Finally, a gender difference in response to tDCS has already been reported in at least one study (Chaieb, Antal, and Paulus 2008).

Studies have shown that DLPFC activity is essential for inhibitory control (IC) (Loftus, Yalcin, Baughman, Vanman, and Hagger 2015) and sustained attention (SA) processes (Plewnia, Zwissler, Längst, et al. 2013; Nelson, McKinley, Golob, et al. 2014). Improvements in response inhibition were observed in studies that applied a-tDCS (using a wide variety of parameters) over l-DLPFC (Jeon and Han 2012; Boggio, Bermpohl, Vergara, et al. 2007). L-DLPFC was reported as important for representing and maintaining the attentional demands of a task in order to implement cognitive control during conflict situations monitored by the anterior cingulate cortex (ACC), hence these brain areas are both essential for successful behavior on the Stroop task (MacDonald, Cohen, Stenger, et al. 2000; Aron, Monsell, Sahakian, and Robbins 2004). Cieslik (Cieslik, Zilles, Caspers, et al. 2012) found a strong connectivity between the anterior subregion of DLPFC and ACC associated with monitoring, attention and action inhibition processes.

In view of this body of evidence, the objective of this study was to assess possible effects of c-tDCS over l-DLPFC on the performance of undergraduate students in the Stroop and Reaction Time tasks. We hypothesized that c-tDCS would have a negative effect on subject performance, i.e., that it would lead to an impairment of IC and attention abilities, since this type of stimulation is presumably associated with reduction of cortical excitability and l-DLPFC plays an essential role in the cognitive tests performed. We predicted that subjects would possibly perform more slowly in the tests and make more errors after c-tDCS when compared to the sham session. We did not attempt to improve performance by means of a-tDCS, since young healthy subjects already perform these tasks at optimal levels (Sá 2015; Canabarro, Garcia, Satler, and Tavares 2017) and this effect of a-tDCS has already been well documented (Dedoncker, Brunoni, Baeken, and Vanderhasselt 2016).

Regarding the possibility of gender differences in inhibitory mechanisms, a review reported that women perform better than men in tasks in the social and behavioral domains, including motor inhibition tasks, but did not find gender differences in performance of cognitive tasks, such as the Stroop test (Bjorklund and Kipp 1996). On the other hand, a gender difference in the effects of neuromodulation by tDCS has also been suggested (Chaieb, Antal, and Paulus 2008). Therefore, this study also aimed at investigating possible gender differences in neuromodulation by c-tDCS.

## Methods

### Participants

Seventeen healthy undergraduate students (ten women) from the University of Brasilia (UnB) served as subjects. Participants ranged in age from 19 to 27 years (mean = 21.4, SE = 0.5) and were recruited by internet advertisements and at the University campus. All subjects were volunteers and signed an informed consent document. The study was approved by the local Ethics Committee. All participants had slept without interruption for at least five hours the night before the experiment, had normal or corrected-to-normal vision and reported no history of personal or family neurological or psychiatric disorders and no medication, alcohol or caffeine intake within the 24 hours prior to the procedure. They were right-handed according to the Edinburgh Handedness Inventory (Oldfield 1971) and had normal color vision according to the Ishihara test. Subjects who did not meet these criteria performed the experiment, but their data were not analyzed.

### Procedure

The study took place in two sessions: a real c-tDCS session,followed by the performance of neuropsychological tests by the subjects, and a sham c-tDCS session, followed by the performance of the same tests. Participants were randomly assigned to one of two groups (Figure 1). The first group performed the real c-tDCS session and returned a week later in order to perform the sham c-tDCS session. The order of the real and sham c-tDCS sessions was reversed for the participants in the second group.

**Figure 1:**
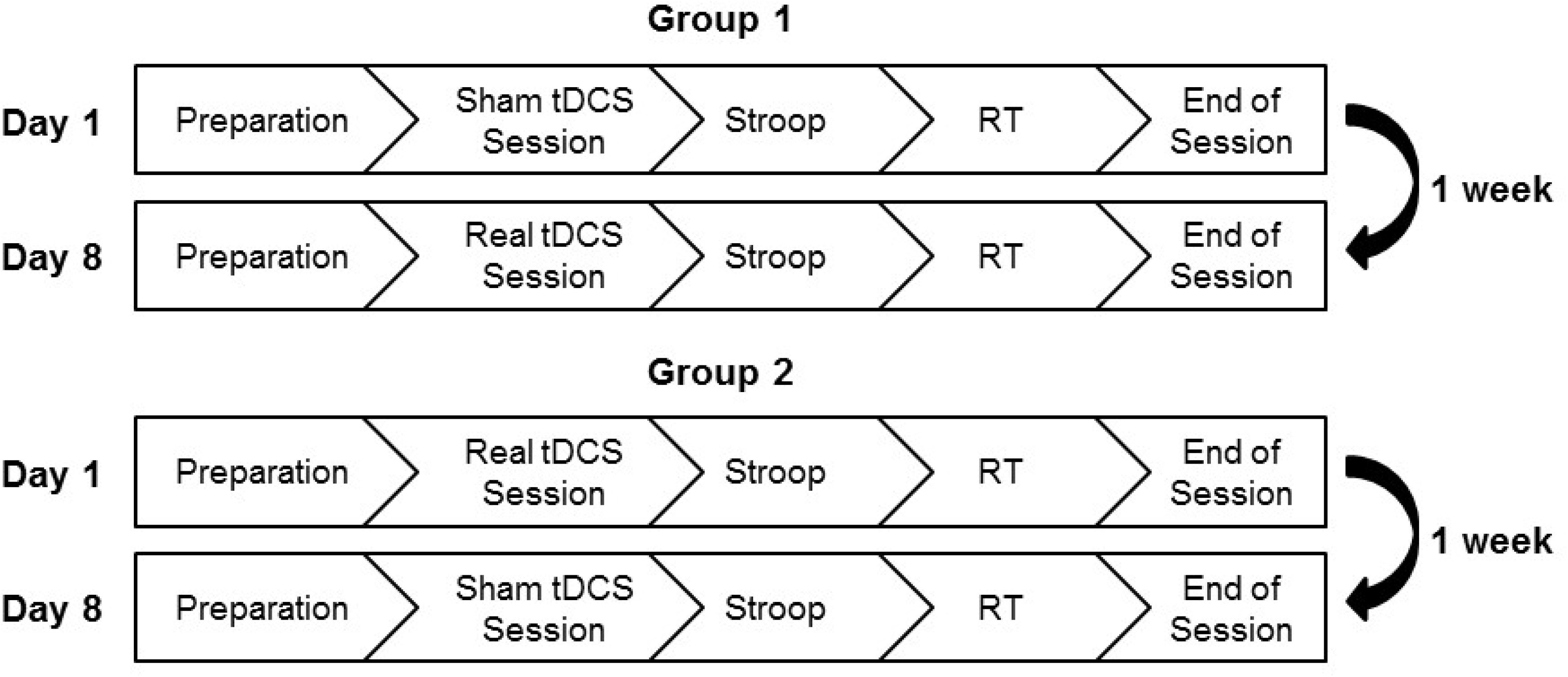
Study procedure. tDCS: cathodal transcranial direct current stimulation, Stroop: Stroop task, RT task: Reaction Time task.

### c-tDCS

C-tDCS was applied using the Trans-Cranial tDCS Stimulator device (Trans-Cranial®, Hong Kong, China) during 20 minutes in two sessions, real and sham, separated by an interval of one week and ordered according to the subject’s group. A pair of sponge electrodes moistened with saline was used. The cathode (25 cm^2^) was positioned on the F3 site of the 10-20 international EEG system, which corresponds to l-DLPFC (Plewnia, Zwissler, Längst, et al. 2013; Jeon and Han 2012; Boggio, Bermpohl, Vergara, et al. 2007). The anode (35 cm^2^) was used as a cephalic reference electrode and was positioned on the contralateral supraorbital area. The large size of such reference electrode renders the stimulation over the contralateral orbitofrontal cortex ineffective (Nitsche and Paulus 2000).

Current intensity gradually increased during 30 seconds in the beginning of the stimulation until it reached 1mA. In the real session, current intensity was maintained at 1mA during 20 minutes, and then gradually decreased at the final seconds of the session. In the sham session, the stimulation was tapered off and interrupted after the first 30 seconds, but the electrodes remained in place during 20 minutes. Subjects were not informed if the session was real or sham.

In order to induce a functional targeting effect, since tDCS seems to affect more strongly neurons that are actively engaged in a task (Villamar, Wivatvongvana, Patumanond, et al. 2013), neuropsychological tests that demand the activity of the DLPFC were performed by the subjects, such as the Flanker task, Iowa Gambling task, Simon task, Oddball task, Spatial cueing task and Psychomotor Vigilance task, all included on the PEBL Test Battery (Mueller and Piper 2014) and translated to Portuguese in our laboratory. These tests were presented in the above mentioned order, as many as the subjects could perform within the 20 minutes of stimulation, as their duration depends on each individual’s performance.

### Stroop task

A computerized version of the Stroop task that belongs to the PEBL Test Battery (Mueller and Piper 2014) adapted in our laboratory was used. One word was presented at a time during 800 ms at the center of a grey background. Participants were instructed to pronounce loudly the color of the words. The task had three stages with 32 trials each. There was an interval of 200 ms between trials. In the first two stages (congruent stage, CS and incongruent stage, IS) all the words were names of colors (red, blue, green and yellow, in Portuguese: *vermelho, azul, verde* and *amarelo*), but in the last stage (phonetic similarity stage, PSS) words were phonetically similar to names of colors (in Portuguese: *velho, cabul, verdade* and *marmelo*). In the first stage, both word attributes (color and the word itself) were congruent (Figure 2), for example the word red written in red. In the last two stages, an interference factor was present as those two attributes could be congruent or not, for example, the word red written in blue. Words were presented in a pseudo-random order. Answers given by the subjects and the computer screen were recorded using a video camera (SMX-C200BN, Samsung®, China).

**Figure 2:**
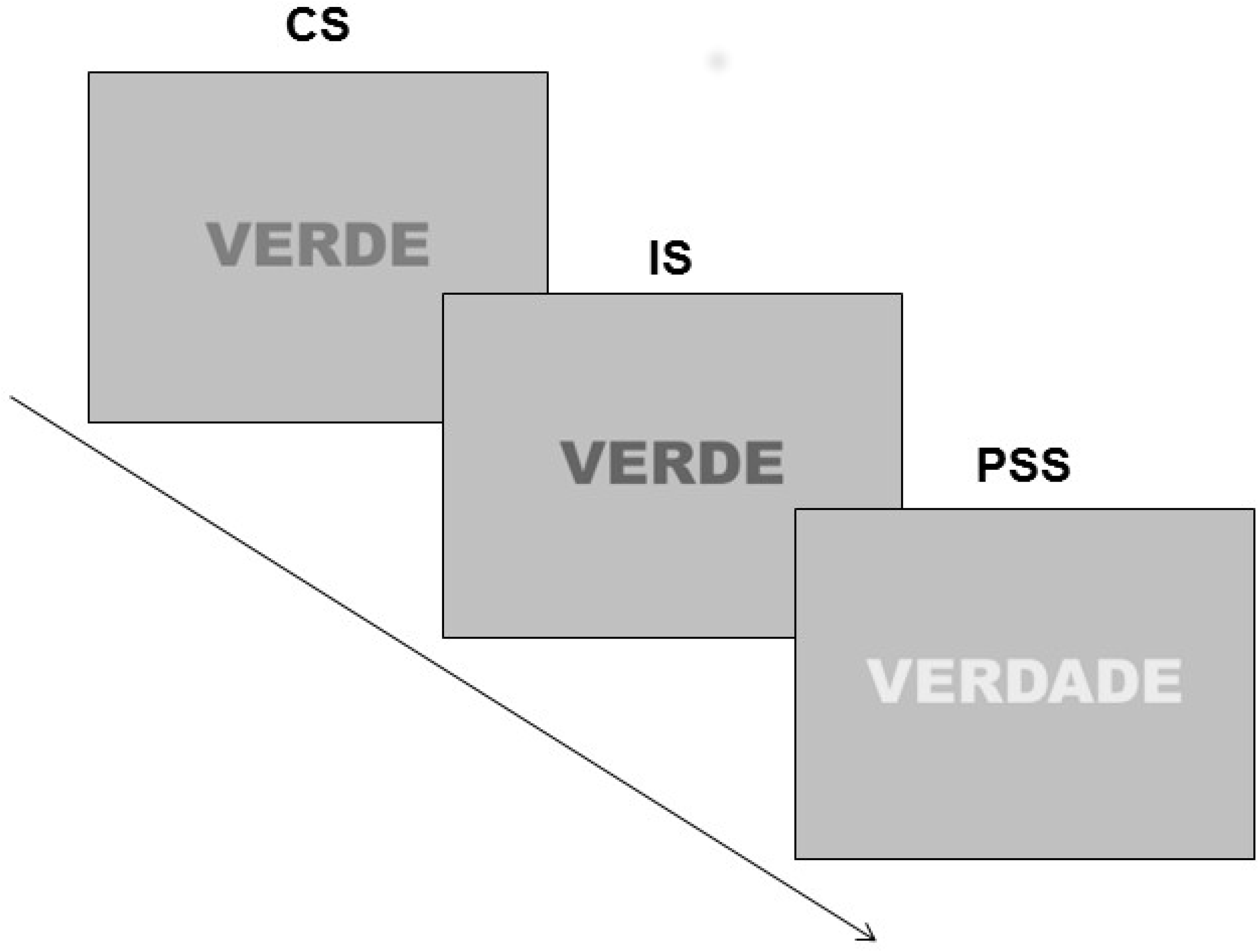
Sequence of the Stroop task with sample images of each stage.

### Reaction Time task

A computerized version of the Reaction Time task (Figure 3) was developed and validated at our laboratory in C language: REFLEX software, v. 1.0. In each trial, a yellow cross (0.02×0.02 m) was shown at the center of a black background and after a random variable interval between 100 and 2000 ms, a target consisting of an unfilled square with white borders (0.03×0.03 m) was presented at one of six possible screen positions (above, in the middle or below, to the left or to the right). Subjects were instructed to answer to the target as fast and as accurately as possible by pressing the space bar on the keyboard. After the response, the target and the cross disappeared and after 1000 ms a new trial began. If there was no response, both stimuli disappeared after 5 s. 150 trials were presented in approximately 6 minutes, depending on the time each subject took to perform each trial.

**Figure 3:**
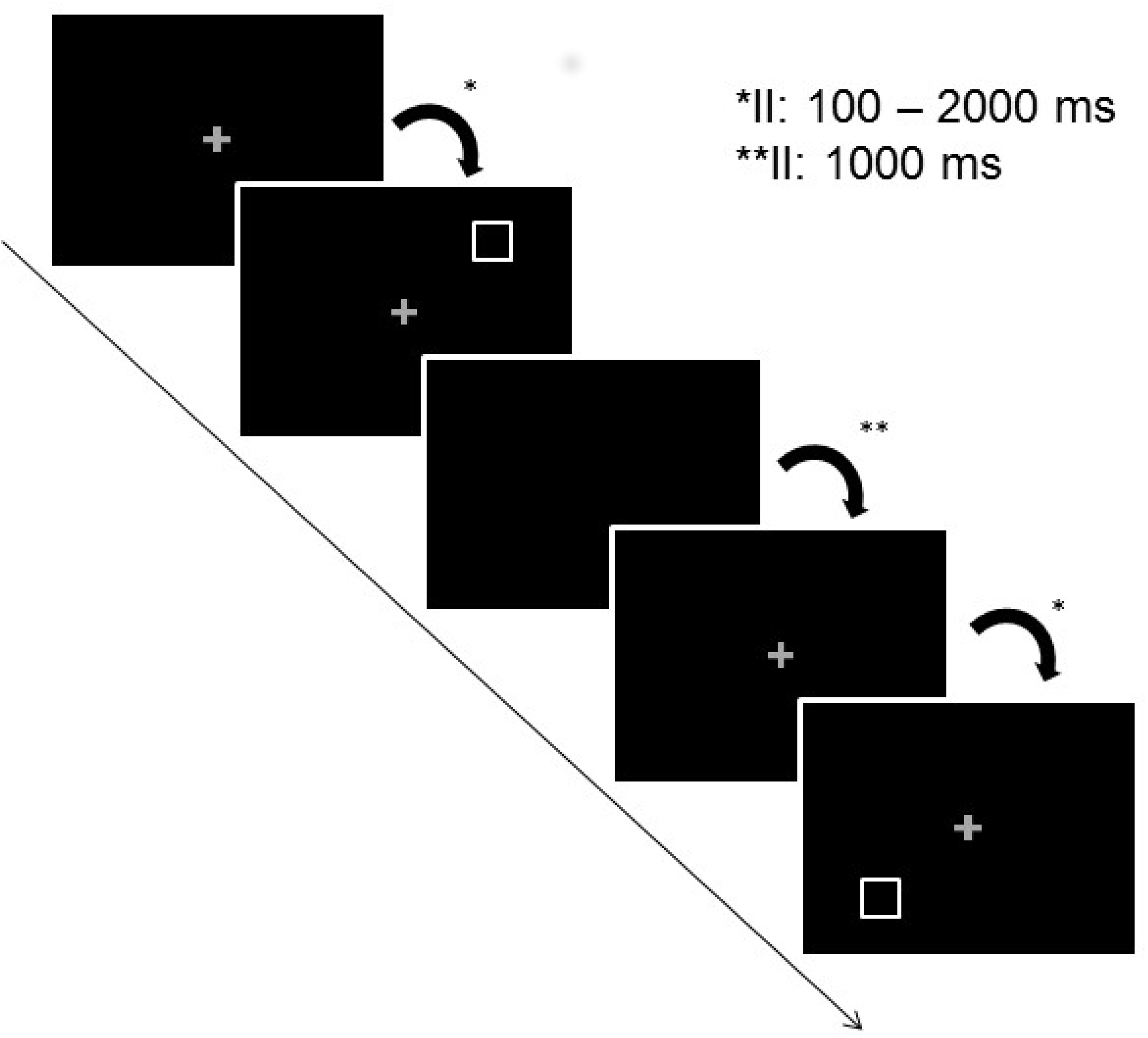
Sequence and sample images of the Reaction Time task. II: Intertrial interval.

## Results

### Stroop task

Behavioral performance parameters for each stage of the Stroop task were measured by the number of hits, errors and omission errors as well as the mean Reaction time of the participants. Trials with no response were considered as omission errors. The Reaction time (ms) for each trial was measured as the elapsed time between the presentation of the word and the onset of the subject’s response.

Two-way repeated measures analysis of variance (ANOVA, Session x Stage, 2×3) performed using the PSAW Statistics software (v. 18.0 for Windows) was used to compare performance parameters between stages. The level of statistical significance was corrected by the Bonferroni method and was set at 5% (p<0.05) for all tests.

An effect of stage was found for all performance parameters (number of hits: F(2,96)=11.428 p<0.001, errors: F(2,96)=6.183 p=0.003, omission errors: F(2,96)=11.517 p<0.001, and RT: F(2,96)=20.185 p<0.001). The number of hits was significantly higher in the congruent stage (CS) when compared to the incongruent (IS, p<0.001), and the phonetic similarity stages (PSS, p=0.021) (Figure 4.1). The number of errors was significantly lower in CS when compared to IS (p=0.004) and PSS (p=0.021) (Figure 4.2). The number of omission errors was significantly higher in IS in comparison with CS (p=0.001) and PSS (p=0.001) (Figure 4.3). Subjects answered faster to CS trials when compared to IS (p<0.001) and PSS (p<0.001) (Figure 4.4).

**Figure 4:**
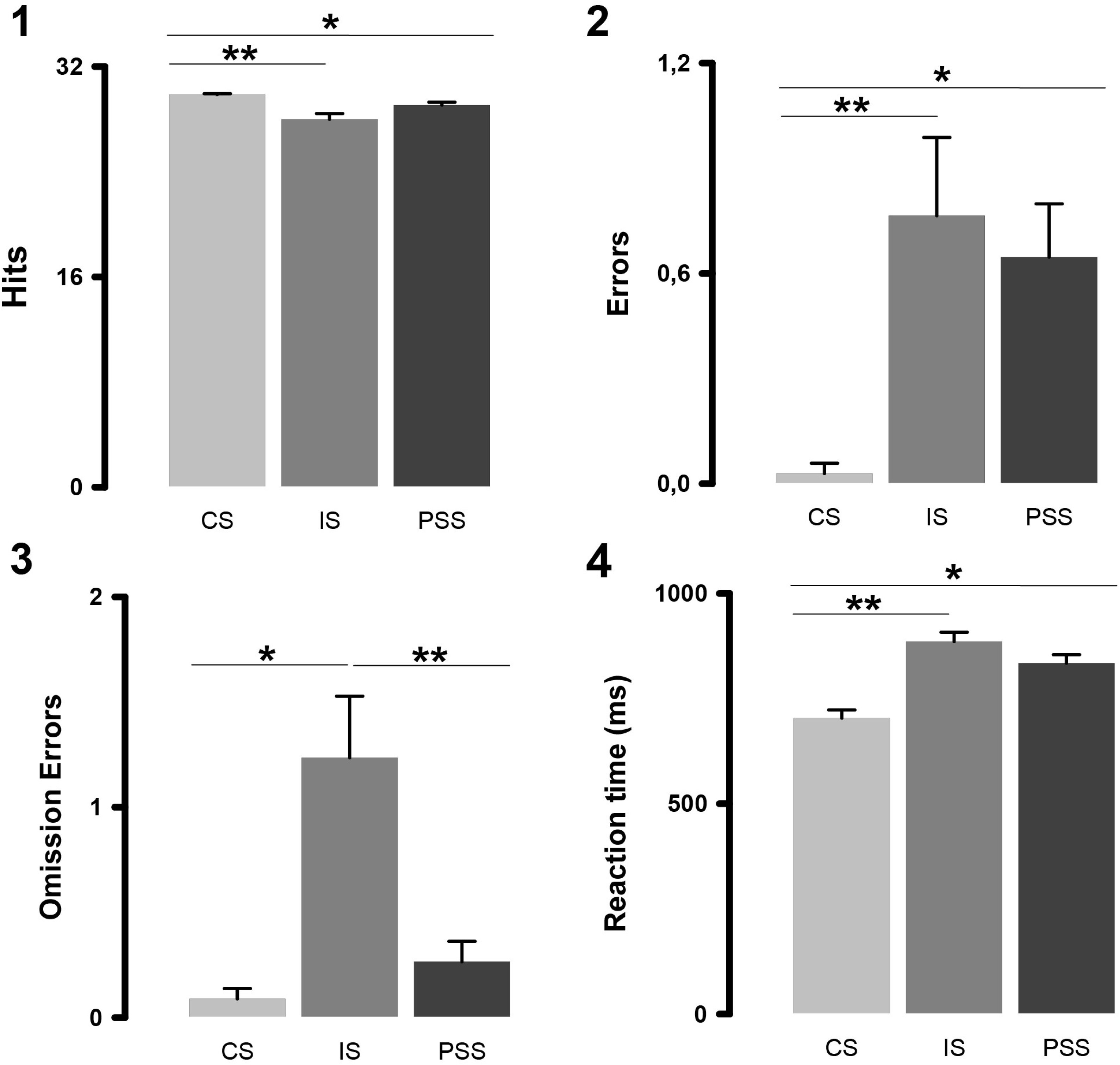
Performance of young undergraduate students (n=17) in each stage of the Stroop task according to the number of hits (1), errors (2), omission errors (3) and reaction time (4). 1: *CS>PSS p<0.05, **CS>IS p<0.001, 2: *CS<PSS p<0.05, **CS<IS p<0.01, 3: *IS>CS p<0.001, **IS>PSS p=0.001, and 4: *CS<PSS p<0.001, **CS<IS p<0.001. CS: congruent stage, IS: incongruent stage and PSS: phonetic similarity stage. The results are presented as mean ± standard error (SE).

Neuromodulation (session) effects were not found for any of the parameters (number of hits: F(1,96)=0.299 p=0.586, errors: F(1,96)=0.286 p=0.594, omission errors: F(1,96)=0.139 p=0.710, and RT: F(1,96)=0.308 p=0.580). Similarly, there was no interaction between session and stage (hits: F(2,96)=0.077 p=0.925, errors: F(2,96)=0.149 p=0.862, omission errors: F(2,96)=0.531 p=0.590, and RT: F(2,96)=0.031 p=0.970).

Considering session results separately, the number of hits in CS was significantly higher than in IS for both real (p=0.002) and sham (p=0.006) sessions (Figure E.1). The number of errors did not differ significantly (Figure 5.2). Both in real (p=0.002) and sham (p=0.022) sessions, the number of omission errors was higher in the incongruent stage in comparison to the congruent stage, but only in the real session an increase in the number of omission errors was found comparing the IS to the phonetic similarity stage (p<0.01) (Figure 5.3). The reaction time was lower in the CS when compared to IS and PSS both in the real (p<0.01 and p<0.001, respectively) and sham (p<0.01 and p<0.001) sessions (Figure 5.4).

**Figure 5:**
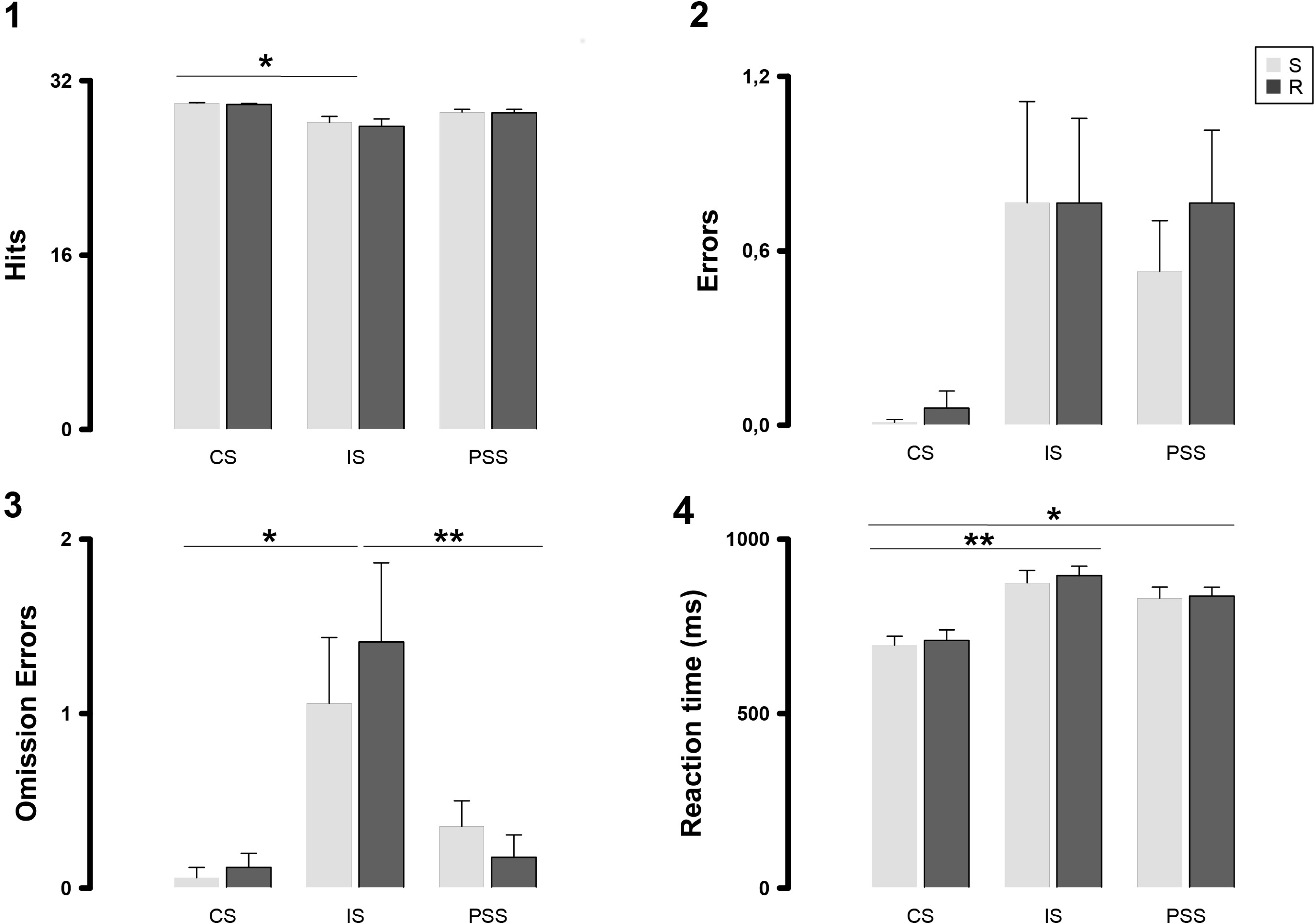
Performance of young undergraduate students (n=17) in each stage of the Stroop task as a function of the neuromodulation session according to the number of hits (1), errors (2), omission errors (3) and reaction time (4). 1 (sham): *CS>IS p<0.01, 1 (real): *CS>IS p<0.01, 3 (sham): *IS<CS p<0.05, 3 (real): *IS>CS p<0.01, **IS>PSS p<0.01, 4 (sham): *CS<PSS p<0.001, **CS<IS p<0.01, and 4 (real): *CS<PSS p<0.001, **CS<IS p<0.01. R: real tDCS session, S: sham tDCS session, CS: congruent stage, IS: incongruent stage, and PSS: phonetic similarity stage. The results are presented as mean ± standard error (SE).

Regarding gender, session effects were not found by the two-way ANOVA (Session x Gender, 2×2) with correction by the Bonferroni method for any of the behavioral parameters (number of hits: F(1,98)=0.207 p=0.650, errors: F(1,98)=0.213 p=0.646, omission errors: F(1,98)=0.098 p=0.755, and RT: F(1,98)=0.221 p=0.640). Gender effects were also not found (number of hits: F(1,98)=0.245 p=0.622, errors: F(1,98)=0.059 p=0.809, omission errors: F(1,98)=0.308 p=0.580, and RT: F(1,98)=0.075 p=0.785), nor were significant interactions found between session and gender (number of hits: F(1,98)=0.037 p=0.849, errors: F(1,98)=0.048 p=0.827, omission errors: F(1,98)=0.012 p=0.912, and RT: F(1,98)=0.002 p=0.969) for any of the parameters.

### Reaction Time task

Behavioral performance on the Reaction Time task was measured by hit rate and reaction time. Answers were considered correct if the response was emitted after target presentation. Errors happened if subjects answered before appearance of the stimuli or immediately after them. The reaction time consisted in the elapsed time (ms) between target presentation and subject response.

Two-way ANOVA (Session x Gender, 2×2) with correction by the Bonferroni method showed no effect of neuromodulation session (hit rate: F(1,30)=0.640 p=0.430, and RT: F(1,30)=0.721 p=0.403), gender (hit rate: F(1,30)=0.003 p=0.960, and RT: F(1,30)=0.119 p=0.733), or of the interaction between session and gender (hit rate: F(1,30)=1,655 p=0.208, and RT: F(1,30)=0.577 p=0.453) on any of the parameters (Table 1).

**Table 1:** Performance of young adults (n=17) on the Reaction Time task as a function of the type of tDCS session (sham or real) considering the hit rate (in percentage) and the reaction time (ms).

| **Behavioral Parameter** | **Measure** | **Real tDCS** | **Sham tDCS** |
| --- | --- | --- | --- |
| Hit rate (%) | Mean (SE) | 98.6 (0.26) | 98.4 (0.25) |
|  | Range | 98.0-99.1 | 97.8-98.9 |
| Reaction time (ms) | Mean (SE) | 349.4 (8.3) | 357.1 (6.2) |
|  | Range | 331.9-367.0 | 344.0-370.2 |

## Discussion

### Stroop Task

Stroop interference effect is defined as an increase in the reaction time when the congruent stage or stimuli are compared to the incongruent ones (Stroop 1935). This is presumably due to the conflict between the word and its color at the response level (Van der Elst, Van Boxtel, Van Breukelen, et al. 2006; Hanslmayr, Pastötter, Bäuml, et al. 2008). For this reason, it was already expected that we would find an effect of stage on the subjects’ performance, that is, we assumed that their reaction time would be longer in the incongruent stage (IS) which could eventually result in a larger number of errors, as seen in a previous work by our group, using the same subjects, but without tDCS (Canabarro, Garcia, Satler, and Tavares 2017).

Our results showed that subjects presented faster reaction times in the congruent stage (CS) compared to the other two stages. Additionally, they made fewer errors in the CS. In the CS, even if subjects read the word, despite this being contrary to the instructions, they did not make an error, since both attributes of the stimuli were congruent. Hence, without a conflict they were able to answer faster and more accurately. However, when they were exposed to the IS, there was a conflict between the two attributes of the word (the color in which it was written and the word itself). In this case, our results showed that subjects took longer to answer to the trials and made more errors than in the CS.

Concerning the number of hits and errors and the reaction time, this pattern was similar to the one found in the phonetic similarity stage (PSS). This indicates that the PSS involved a conflict between the two attributes of the word; however, in this stage there was no incongruity, that is, since the word presented was not the name of a color, those two attributes could not be incongruent in the same sense as they were in the IS. Thus, for both IS and PSS at the response level there was likely a competition between the two possibilities of response (there was a conflict), but for the former, those two possibilities were incongruent. This may help to explain why the number of omission errors was higher in the IS when compared to the CS and PSS, but did not significantly differ between these last two.

We assumed that the omission errors happened when the difficulty in one trial was so high that it impaired response in subsequent trials. In a stage where there is not only conflict but also incongruity as the IS, it is expected that the number of omission errors increases, as it was in fact observed.

We have thus regarded PSS as a situation of intermediate difficulty, being more difficult than the CS where there was no conflict at all, but easier than the IS where there was conflict and also incongruity. Other studies that included stages with neutral words in the same task, i. e, words that were not names of colors as in our PSS, also found higher reaction times than in congruent stages, but lower than in incongruent ones (Hanslmayr, Pastötter, Bäuml, et al. 2008; Mead, Mayer, Bobholz, et al. 2002). These results can be interpreted on the basis of inhibitory control. The demands for inhibiting the automatic action of reading the words instead of naming their colors were not as high in the CS as they were in the IS and PSS.

Regarding c-tDCS, the type of neuromodulation session (sham or real) did not have any effect on behavioral performance. An interesting result was found considering performance in the two sessions separately. The results showed that the number of omission errors was higher in IS in comparison with CS in both real and sham c-tDCS sessions. However, only in real sessions was the number of omission errors in IS significantly higher than in PSS. This could suggest an impairment in behavioral performance during the IS after delivery of c-tDCS to l-DLPFC. However, due to the lack of the main effect of c-tDCS session on performance according to the statistical analysis, we assumed that this result is probably an error (type I) generated by the large number of statistical tests that were run.

Loftus (Loftus, Yalcin, Baughman, Vanman, and Hagger 2015) conducted a study in which participants performed the Stroop task before and after sham or a-tDCS over l-DLPFC (in the real session 1 mA was applied during 10 min). As in our study, their results showed that raw reaction times did not significantly differ between anodal and sham a-tDCS. However, they found significant a-tDCS effects on the reaction time “change scores” (subtraction between post-tDCS and pre-tDCS Stroop RT), that is, subjects were faster after stimulation of l-DLPFC. These results highlight the importance of new studies on the different effects of a-tDCS and c-tDCS upon cognitive function.

Gender did not influence behavioral performance in any of the measured parameters. This goes in line with the hypothesis developed by Bjorklund and Kipp (Bjorklund and Kipp 1996) of no marked differences between the genders in tasks involving IC in the cognitive domain. Similarly, there was no differential effect of c-tDCS between genders, which indicates a similar lack of effect of c-tDCS applied over l-DLPFC for both men and women.

### Reaction Time task

The hit rate in the Reaction Time task was high in every study condition. The occurrence of a ceiling effect in the behavioral performance can be explained by the fact that participants were normal young adults with no impairments in processing speed or in sustained attention. However, reaction times were relatively prolonged in comparison with those found in other studies involving similar tasks (Thut, Hauert, Morand, Seeck, Landis, and Michel 1999; Barbarotto, Laiacona, Frosio, Vecchio, Farinato, and Capitani 1998; Serrien, Fisher, and Brown 2003). We assume that we used a more complex version of the RT task and hence, subjects took longer to respond to the trials, but kept response accuracy.These results are in line with a study by our group that used a version of the same task with less trials (72) in the same population (Sá 2015).Apparently for this population the increased number of trials in this version (150) was still not enough to prevent the ceiling effect.

Delivery of c-tDCS over l-DLPFC had no effect on behavioral performance in this task. In our study, the ceiling effect may have hampered the assessment of possible beneficial c-tDCS effects on the hit rate. However, a study assessing SA and response inhibition during a Go/No-Go task (Boggio, Bermpohl, Vergara, et al. 2007) did not find any effects of a-tDCS (20 min, 1mA) over the DLPFC on the hit rate. Therefore, for future studies, it is advisable to use a longer version of the RT task, including more trials or increasing the inter-trial interval, in order to clarify this issue by reducing sustained attention due to vigilance decrement effects.

We did not find gender differences either in the performance in the RT task itself or in c-tDCS effects. In the literature, it is not clear whether there are gender differences in the RT task. While Bjorklund (Bjorklund and Kipp 1996) found that women perform better that men in motor inhibition tasks, Der and Deary (Der and Deary 2006) claim that, when found, gender differences in reaction times are due to performance variability during a task owing to a trade-off between speed and accuracy in the same subject. Therefore, in their study, women were slower and more accurate than men in the beginning of the task, but became faster and kept their accuracy over time.

## Conclusion

From this work, several conclusions can be drawn:

1. Our behavioral findings are consistent with previous studies showing an expected interference effect during the Stroop task. This was different in the RT task, for which we found a ceiling effect.
2. Regarding c-tDCS over l-DLPFC, there were no significant effects (positive or negative) upon performance of the Stroop or the Reaction Time tasks. These results are clearly different from what has been described in the literature for a-tDCS (Nitsche and Paulus 2000; Jeon and Han 2012; Hsu, Tseng, Yu, et al. 2011; Jacobson, Javitt, and Lavidor 2011; Ditye, Jacobson, Walsh, et al. 2012; Boggio, Bermpohl, Vergara, et al. 2007; Beeli, Casutt, Baumgartner, et al. 2008).
3. Both male and female participants exhibited similar performance patterns in the tests after a real or sham c-tDCS session, suggesting lack of gender differences as well.

Our results add support to a recent review (Dedoncker, Brunoni, Baeken, and Vanderhasselt 2016) which pointed out a paucity of evidence for c-tDCS effects upon cognitive function.

Likewise, they draw attention to the fact that the excitatory effects of a-tDCS may not necessarily be reversed when c-tDCS is used instead (Jacobson, Javitt, and Lavidor 2011). Caution should therefore be exercised, in future studies in the experimental and clinical realms, when assuming that c-tDCS would have an inhibitory effect upon any cortical region, as it does in the motor cortex (Nitsche and Paulus 2000).

Additional studies on the effects of c-tDCS upon cognitive function are warranted, since recent research which simultaneously stimulates both prefrontal cortices (Leite, Carvalho, Fregni, Boggio, and Gonçalves 2013; Loftus, Yalcin, Baughman, Vanman, and Hagger 2015) is not capable of shedding light on the role of c-tDCS proper.

## Acknowledgements

This research was supported by a CAPES (Brazilian Federal Agency for Support and Evaluation of Graduate Education, Ministry of Education, Brazil) scholarship to S. L. S. Canabarro as well as CNPq (National Counsel of Technological and Scientific Development, Ministry of Science, Technology and Innovation, Brazil) scholarships to C. K. Paniago and P. M. Santos.

## References

Aron, A. R., S. Monsell, B. J. Sahakian, and T. W. Robbins. 2004. “A componential analysis of task-switching deficits associated with lesions of left and right frontal cortex.” Brain 127:1561–1573.

Barbarotto, R, M Laiacona, R Frosio, M Vecchio, A Farinato, and E Capitani. 1998. “A normative study on visual reaction times and two Stroop colour-word tests.” The Italian Journal of Neurological Sciences 19:161–170.

Beeli, G., G. Casutt, T. Baumgartner, et al. 2008. “Modulating presence and impulsiveness by external stimulation of the brain.” Behavioral and Brain Functions 4:33.

Bjorklund, D.F. and K. Kipp. 1996. “Parental investment theory and gender differences in the evolution of inhibition mechanisms.” Psychological bulletin 120:163.

Boggio, P.S., F. Bermpohl, A.O. Vergara, et al. 2007. “Go-no-go task performance improvement after anodal transcranial DC stimulation of the left dorsolateral prefrontal cortex in major depression.” Journal of affective disorders 101:91–98.

Canabarro, SL, A Garcia, C Satler, and MCH Tavares. 2017. “Interaction between Neural and Cardiac Systems during the Execution of the Stroop Task by Young Adults: Electroencephalographic Activity and Heart Rate Variability.” AIMS Neuroscience 4:28–51.

Chaieb, L., A. Antal, and W. Paulus. 2008. “Gender-specific modulation of short-term neuroplasticity in the visual cortex induced by transcranial direct current stimulation.” Visual neuroscience 25:77–81.

Cieslik, E.C., K. Zilles, S. Caspers, et al. 2012. “Is there “one” DLPFC in cognitive action control? Evidence for heterogeneity from co-activation-based parcellation.” Cerebral cortex p. bhs256.

Dedoncker, Josefien, Andre R. Brunoni, Chris Baeken, and Marie-Anne Vanderhasselt. 2016. “A Systematic Review and Meta-Analysis of the Effects of Transcranial Direct Current Stimulation (tDCS) Over the Dorsolateral Prefrontal Cortex in Healthy and Neuropsychiatric Samples: Influence of Stimulation Parameters.” Brain Stimulation 9:501–517.

Der, G. and I.J. Deary. 2006. “Age and sex differences in reaction time in adulthood: results from the United Kingdom Health and Lifestyle Survey.” Psychology and aging 21:62.

Ditye, T., L. Jacobson, V. Walsh, et al. 2012. “Modulating behavioral inhibition by tDCS combined with cognitive training.” Experimental brain research 219:363–368.

Filmer, Hannah L., Jason B. Mattingley, and Paul E. Dux. 2013. “Improved multitasking following prefrontal tDCS.” Cortex 49:2845–2852.

Hanslmayr, S., B. Pastötter, K. Bäuml, et al. 2008. “The electrophysiological dynamics of interference during the Stroop task.” Journal of Cognitive Neuroscience 20:215–225.

Hsu, T., L. Tseng, J. Yu, et al. 2011. “Modulating inhibitory control with direct current stimulation of the superior medial frontal cortex.” Neuroimage 56:2249–2257.

Jacobson, L., D.C. Javitt, and M. Lavidor. 2011. “Activation of inhibition: diminishing impulsive behavior by direct current stimulation over the inferior frontal gyrus.” Journal of Cognitive Neuroscience 23:3380–3387.

Jeon, S.Y. and S.J. Han. 2012. “Improvement of the working memory and naming by transcranial direct current stimulation.” Annals of rehabilitation medicine 36:585–595.

Kang, E., D. Kim, and N. Paik. 2012. “Transcranial direct current stimulation of the left prefrontal cortex improves attention in patients with traumatic brain injury: a pilot study.” Journal of rehabilitation medicine 44:346–350.

Kwon, Y. H. and J. W. Kwon. 2013. “Is transcranial direct current stimulation a potential method for improving response inhibition?” Neural Regeneration Research 8:1048–1054.

Leite, Jorge, Sandra Carvalho, Felipe Fregni, Paulo S Boggio, and Óscar F Gonçalves. 2013. “The effects of cross-hemispheric dorsolateral prefrontal cortex transcranial direct current stimulation (tDCS) on task switching.” Brain stimulation 6:660–667.

Leite, Jorge, Sandra Carvalho, Felipe Fregni, and Óscar F Gonçalves. 2011. “Task-specific effects of tDCS-induced cortical excitability changes on cognitive and motor sequence set shifting performance.” PloS one 6:e24140.

Loftus, A.M., O. Yalcin, F.D. Baughman, E.J. Vanman, and M.S. Hagger. 2015. “The impact of transcranial direct current stimulation on inhibitory control in young adults.” Brain and behavior 5.

MacDonald, A.W., J.D. Cohen, V.A. Stenger, et al. 2000. “Dissociating the role of the dorsolateral prefrontal and anterior cingulate cortex in cognitive control.” Science 288:1835–1838.

Mead, L.A., A.R. Mayer, J.A. Bobholz, et al. 2002. “Neural basis of the Stroop interference task: response competition or selective attention?” Journal of the International Neuropsychological Society 8:735–742.

Miniussi, C. and M. Ruzzoli. 2012. “Transcranial stimulation and cognition.” Handbook of clinical neurology 116:739–750.

Mueller, S.T. and B.J. Piper. 2014. “The psychology experiment building language (PEBL) and PEBL test battery.” Journal of neuroscience methods 222:250–259.

Nelson, J.T., R.A. McKinley, E.J. Golob, et al. 2014. “Enhancing vigilance in operators with prefrontal cortex transcranial direct current stimulation (tDCS).” Neuroimage 85:909–917.

Nitsche, M.A., L.G. Cohen, E.M Wassermann, et al. 2008. “Transcranial direct current stimulation: state of the art 2008.” Brain stimulation 1:206–223.

Nitsche, M. A and W. Paulus. 2000. “Excitability changes induced in the human motor cortex by weak transcranial direct current stimulation.” The Journal of physiology 527:633–639.

Oldfield, Richard C. 1971. “The assessment and analysis of handedness: the Edinburgh inventory.” Neuropsychologia 9:97–113.

Paulus, W., A. Antal, and M.A. Nitsche. 2013. “Physiological Basis and Methodological Aspects of Transcranial Electric Stimulation (tDCS, tACS and tRNS).” In Transcranial Brain Stimulation, edited by C. Miniussi, W. Paulus, and P.M. Rossini, pp. 93–111. Florida: CRC Press.

Plewnia, C., B. Zwissler, I. Längst, et al. 2013. “Effects of transcranial direct current stimulation (tDCS) on executive functions: influence of COMT Val/Met polymorphism.” Cortex 49:1801–1807.

Sarkis, R.A., N. Kaur, and J.A. Camprodon. 2014. “Transcranial direct current stimulation (tDCS): modulation of executive function in health and disease.” Current Behavioral Neuroscience Reports 1:74–85.

Serrien, D.J., R.J. Fisher, and P. Brown. 2003. “Transient increases of synchronized neural activity during movement preparation: influence of cognitive constraints.” Experimental brain research 153:27–34.

Stroop, J.R. 1935. “Studies of interference in serial verbal reactions.” Journal of experimental psychology 18:643.

Sá, S.L. 2015. Behavioral and Electroencephalographic Analysis of Motor and Verbal Inhibitory Control in Young University Students. Master’s thesis, University of Brasília, Brasília, Brazil.

Thut, Gregor, Claude-Alain Hauert, Stéphanie Morand, Margitta Seeck, Theodor Landis, and Christoph Michel. 1999. “Evidence for interhemispheric motor-level transfer in a simple reaction time task: an EEG study.” Experimental Brain Research 128:256–261.

Van der Elst, W., M.P.J. Van Boxtel, G.J.P. Van Breukelen, et al. 2006. “The Stroop Color-Word Test influence of age, sex, and education; and normative data for a large sample across the adult age range.” Assessment 13:62–79.

Villamar, M. F., P. Wivatvongvana, J. Patumanond, et al. 2013. “Focal modulation of the primary motor cortex in fibromyalgia using 4×1-ring high-definition transcranial direct current stimulation (HD-tDCS): immediate and delayed analgesic effects of cathodal and anodal stimulation.” J Pain 14:371–383.

